# Antimicrobial mitochondrial reactive oxygen species induction by lung epithelial metabolic reprogramming

**DOI:** 10.1101/2023.01.19.524841

**Authors:** Yongxing Wang, Vikram V. Kulkarni, Jezreel Pantaleón García, Miguel M. Leiva-Juárez, David L. Goldblatt, Fahad Gulraiz, Jichao Chen, Sri Ramya Donepudi, Philip L. Lorenzi, Hao Wang, Lee-Jun Wong, Michael J. Tuvim, Scott E. Evans

## Abstract

Pneumonia is a worldwide threat, making discovery of novel means to combat lower respiratory tract infections an urgent need. We have previously shown that manipulating the lungs’ intrinsic host defenses by therapeutic delivery of a unique dyad of pathogen-associated molecular patterns protects mice against pneumonia in a reactive oxygen species (ROS)-dependent manner. Here we show that antimicrobial ROS are induced from lung epithelial cells by interactions of CpG oligodeoxynucleotides (ODNs) with mitochondrial voltage-dependent anion channel 1 (VDAC1) without dependence on Toll-like receptor 9 (TLR9). The ODN-VDAC1 interaction alters cellular ATP/ADP/AMP localization, increases delivery of electrons to the electron transport chain (ETC), enhances mitochondrial membrane potential (Δ_Ψm_), and differentially modulates ETC complex activities. These combined effects promote leak of electrons from ETC complex III, resulting in superoxide formation. The ODN-induced mitochondrial ROS yield protective antibacterial effects. Together, these studies identify a therapeutic metabolic manipulation strategy that has the potential to broadly protect patients against pneumonia during periods of peak vulnerability without reliance on currently available antibiotics.

**Author Summary:** Pneumonia is a major cause of death worldwide. Increasing antibiotic resistance and expanding immunocompromised populations continue to enhance the clinical urgency to find new strategies to prevent and treat pneumonia. We have identified a novel inhaled therapeutic that stimulates lung epithelial defenses to protect mice against pneumonia in a manner that depends on production of reactive oxygen species (ROS). Here, we report that the induction of protective ROS from lung epithelial mitochondria occurs following the interaction of one component of the treatment, an oligodeoxynucleotide, with the mitochondrial voltage-dependent anion channel 1. This interaction alters energy transfer between the mitochondria and the cytosol, resulting in metabolic reprogramming that drives more electrons into the electron transport chain, then causes electrons to leak from the electron transport chain to form protective ROS. While antioxidant therapies are endorsed in many other disease states, we present here an example of therapeutic induction of ROS that is associated with broad protection against pneumonia without reliance on administration of antibiotics.

## INTRODUCTION

Pneumonia has long been recognized as a leading cause of death among healthy and immunosuppressed people worldwide (1-3). Pneumonia management has historically focused on patient-extrinsic factors, such as antibiotic administration (4, 5). To address such challenges as increasing antibiotic resistance and newly emerging infections, our laboratory focuses on manipulating vulnerable patients’ intrinsic antimicrobial defenses to broadly protect them against pneumonia. We advance a strategy of activating the lungs’ mucosal defenses to induce broad, pathogen-agnostic protection via airway delivery of synthetic Toll-like receptor (TLR) agonists.

Once regarded as simple airflow conduits or inert gas exchange barriers, the airway and alveolar epithelia are critical immune effector cells that supplement the lungs’ mucosal immune defenses by undergoing fundamental structural and functional changes upon encountering pathogens (6-8). These cells sense pathogens via pattern recognition receptors (PRRs), modulate lung leukocyte responses through cytokine and chemokine expression, and release microbicidal molecules such as reactive oxygen species (ROS) and antimicrobial polypeptides (AMPs) (9-11). Harnessing this defensive immune function, we developed a protective PRR agonist therapeutic comprised of a synthetic diacylated lipopetide ligand for TLR2/6 (Pam2CSK4, “Pam2”) and a class C unmethylated CpG oligodeoxynucleotide ligand for TLR9 (ODN M362, “ODN”). A single inhaled treatment with this non-intuitive dyad of ligands (“Pam2-ODN”) for spatially segregated TLRs yields substantial protection against pneumonia (12-16).

We recently reported that Pam2-ODN-induced antimicrobial protection requires therapeutic induction of ROS from both mitochondrial and dual oxidase sources (17, 18), but the molecular mechanisms responsible for inducible antimicrobial ROS generation remained unresolved. Here, we find that ODN induces mitochondrial ROS (mtROS) production via metabolic reprogramming that alters mitochondrial electron transport chain (ETC) activity in a mitochondrial membrane potential (Δ_Ψm_)-dependent manner. These findings provide novel insights into development of metabolic strategies to protect against otherwise lethal pneumonias in vulnerable populations.

## RESULTS

### Induction of epithelial mtROS by CpG ODN

Having reported that Pam2 and ODN M362 are both required for maximal antimicrobial protection in a manner that depends on inducible ROS production from both mitochondria and dual oxidases, we sought to determine which ligand(s) induce mtROS production. As shown in Figure 1A, ODN M362 alone induced as much mtROS generation as the Pam2-ODN combination from human lung epithelial (HBEC3-KT) cells, revealing ODN M362 as the main driver of this response. This capacity to induce mtROS was not common to all nucleic acid treatments (Figure 1A), but was observed following treatment of HBEC-3KT cells with various types of oligodeoxynucleotides (Figure 1B). Similarly, ODN M362 mtROS in mouse lung epithelial (MLE-15) cells (Figure 1C), as well as primary human and mouse lung epithelial cells (Figure 1D-E), regardless of which mtROS detector was used (Figure S1). Fluorescence microscopy of primary lung epithelial cells from mice expressing redox-sensitive mitochondrial GFP (mt-roGFP) (19, 20) revealed that, at baseline, mitochondria display a predominantly reduced GFP phenotype, whereas treatment with ODN induces a predominantly oxidized mitochondrial phenotype (Figure 1F-G). Under normal conditions, oxidative phosphorylation consumes >95% of cellular oxygen (21), but mtROS formation also requires free oxygen (22). By inhibiting oxidative phosphorylation with oligomycin, we demonstrate increased oxygen consumption by superoxide (O_2_-•) production following ODN (Figure 1H). This ODN-induced mtROS production is consistently associated with increased Δ_Ψm_, as assessed by different assays (Figure 1I-K).

**Fig. 1.**
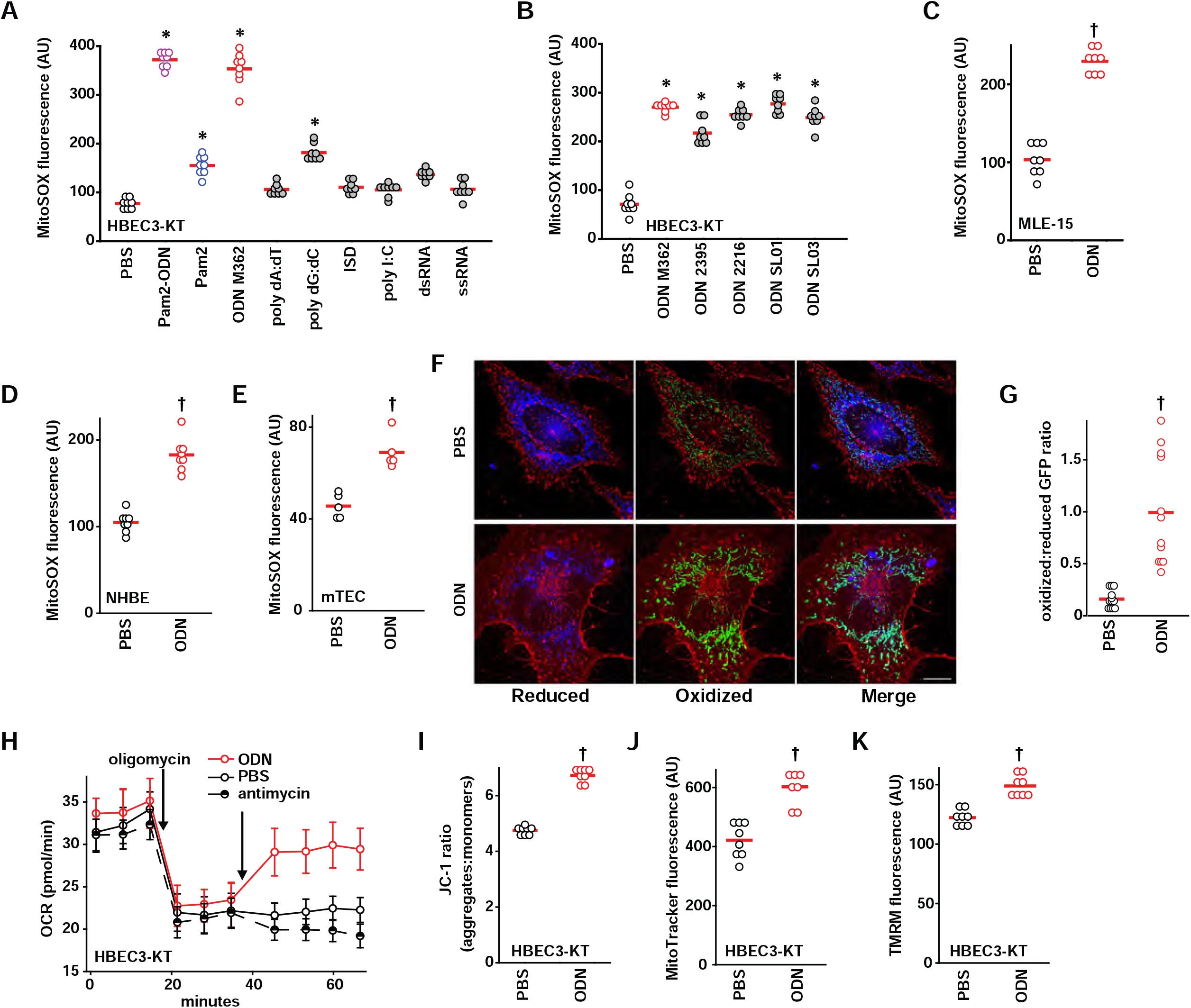
Induction of epithelial mtROS by CpG ODN. (**A**) mtROS production from HBEC3-KT cells after treatment with pathogen associated molecular patterns. (**B**) mtROS production from HBEC3-KT cells after treatment with the indicated ODNs. mtROS production after treatment with ODN from mouse lung epithelial cell lines (**C**) and primary human (**D**) and primary mouse (**E**) lung epithelial cells. (**F**) Representative fluorescence images primary tracheal epithelial cells harvested from mt-roGFP mice treated with PBS or ODN. Images shown as gradient of color intensity from the reduced (blue) form to the oxidized (green) form of roGFP. Scale bar, 50 μm. (**G**) Ratio of the fluorescence intensity of the oxidized:reduced roGFP from **F**, quantified at 488 nm and 405 nm, respectively. (**H**) Oxygen consumption following the indicated treatment by Seahorse XFe96 Flux Analyzer, shown as mean ± SEM. (**I**) Mitochondrial membrane potential Δ_Ψm_ measurement in HBEC3-KT cells after ODN treatment. * p<0.001 vs. PBS by one-way ANOVA using Holm-Sidak method, except A which use Tukey method due to failed normality testing; † p<0.001 vs PBS by two-way Student’s t test. ODN, oligodeoxynucleotide; ISD, immune stimulating DNA; mTEC, primary mouse tracheal epithelial cells; NHBE, primary normal human bronchial epithelial cells; GFP, green fluorescent protein; OCR, oxygen consumption rate; TMRM, tetramethylrhodamine.

### CpG ODN alters electron transport chain activity and energy production

mtROS production is tightly regulated by electron transport chain (ETC) activity (23). To understand the mechanisms of ODN-induced mtROS production, we analyzed the enzymatic activity of the ETC complexes in HBEC3-KT cells, finding that that ODN treatment induces a 35% increase in complex II activity and an 82% decrease in complex III activity, along with an increase in citrate synthase activity (Figure 2A, Figure S2). We also found modest but statistically significant reductions in complex V activity, suggesting ODN may interfere with mitochondrial energy production (Figure S2). The change in ETC complex activity was not accompanied by changes in mitochondrial protein concentrations (Figure 2B, Figure S3), suggesting that the ODN effect is mediated by manipulating ETC function rather than altering mitochondrial mass.

**Fig. 2.**
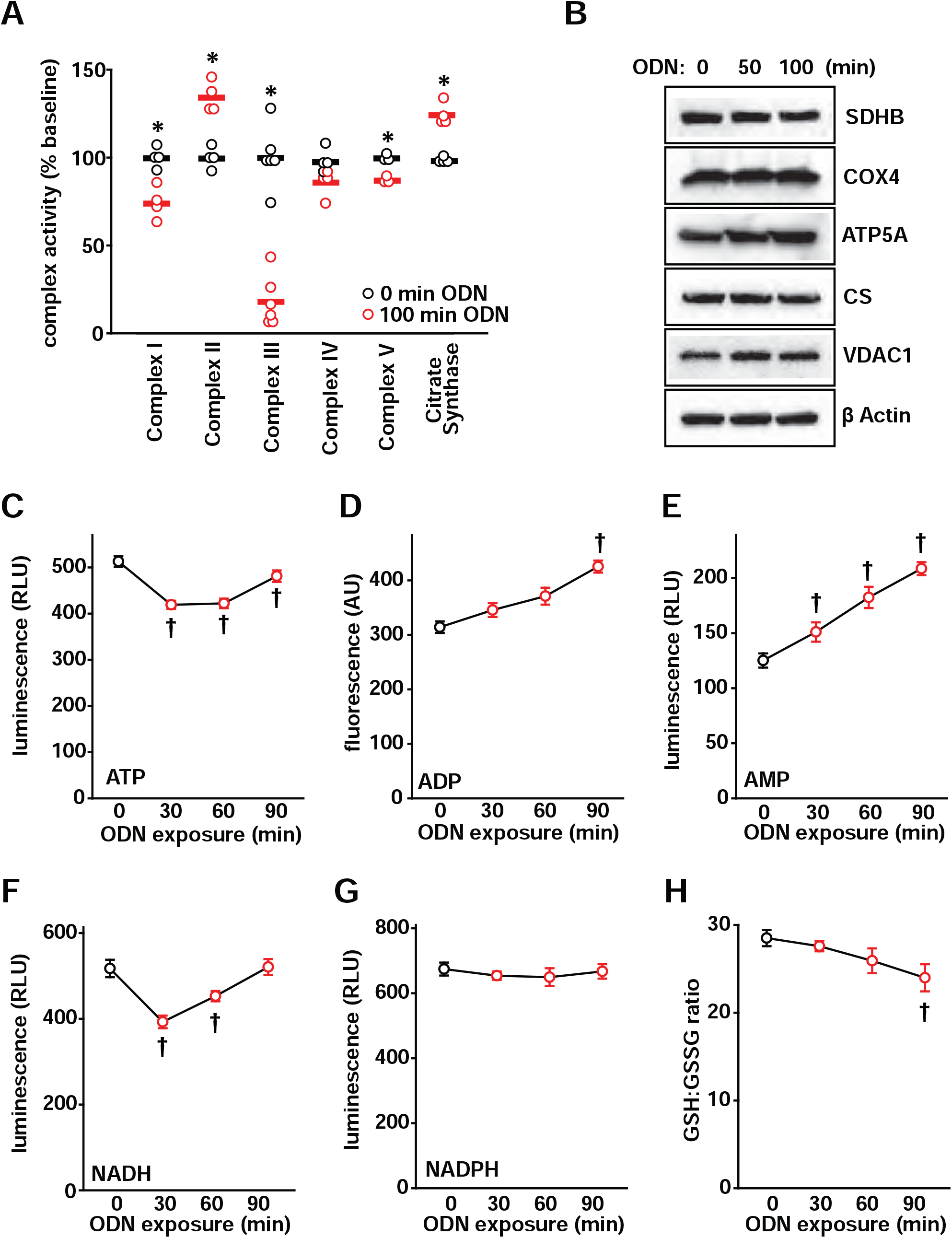
Mitochondrial energy metabolism altered by CpG ODN. (**A**) Summary electron transport chain complex enzyme activity with ODN treatment of HBEC3-KT cells. (**B**) Mitochondrial protein immunoblots from lysates of cells treated with ODN. Relative abundance ATP (**C**), ADP (**D**), and AMP (**E**) in whole cell lysates at the indicated time points after ODN treatment. Relative abundance of NADH (**F**) an NADPH (**G**) in whole cells after ODN treatment. (**H**) Ratio of reduced:oxidized glutathione in HBEC3-KT cells treated with ODN. * p<0.003 vs PBS; † p<0.001 vs PBS. SDHB, succinate dehydrogenase subunit B; COX4, cytochrome c oxidase subunit IV; ATP5A, ATP synthase subunit alpha; CS, citrate synthase; VDAC1, voltage dependent anion channel 1; GSH, reduced glutathione; GSSG, oxidized glutathione.

Since the major energy output of ETC activity is ATP, we investigated the impact of ODN treatment on cellular ATP levels. ODN treatment caused a rapid decline in whole-cell ATP concentrations with a nadir around 30 min that recovered by 90 min (Figure 2C). In contrast, cellular ADP and AMP levels persistently rose following ODN exposure (Figure 2D-E). These effects on ATP, ADP and AMP were ODN dose-dependent (Figure S4). A drop-and-recovery pattern similar to that of ATP was seen in cellular NADH levels after ODN treatment, whereas NADPH levels were largely unaffected by ODN treatment (Figure 2F-G, Figure S4). NAD/NADH are electron receptor and donor that links large molecule catabolism to mitochondrial energy production. The congruent temporal patterns of NADH and ATP support a hypothesis that ODN stimulates catabolic reactions related to mitochondrial energy production. In contrast, NADPH is primarily produced in the anabolic pentose phosphate pathway, which ODN does not appear to perturb. Levels of reduced glutathione persistently declined after ODN treatment (Figure 2H), consistent with the continuing production of ODN-induced ROS.

### CpG ODN blocks mitochondrial nucleotide transition

As TLR9 is the established intracellular sensor for CpG ODNs (24), we examined whether TLR9 activation regulates ODN-induced mtROS generation. To our surprise, ODN treatment still fully induced mtROS production in primary mouse lung epithelial cells isolated from *Tlr9* knockout mice or from mice lacking downstream TLR signaling molecules MyD88 or TRAF6 (Figure S5), indicating that ODN-induced mtROS generation does not require TLR9 signaling.

Seeking to identify a TLR9-independent mechanism by which ODN alters mitochondrial energy metabolism, we investigated whether ODNs could directly stimulate mtROS production in isolated mitochondria. Remarkably, we found that direct ODN treatment of mitochondria isolated from HBEC3-KT cells recapitulates the inducible mtROS generation and increased ΔΨm observed in whole cells (Figure 3A-B). Alternately, to examine whether ODN interacts with mitochondria in intact cells, whole cells were treated with fluorescently-labeled ODN, then the mitochondria were isolated and assessed for fluorescence intensity. As in Figure 3C, mitochondria from cells treated with labeled ODN displayed significant fluorescence, supporting a hypothesis that ODN can directly interact with mitochondria to stimulate mtROS production.

**Fig. 3.**
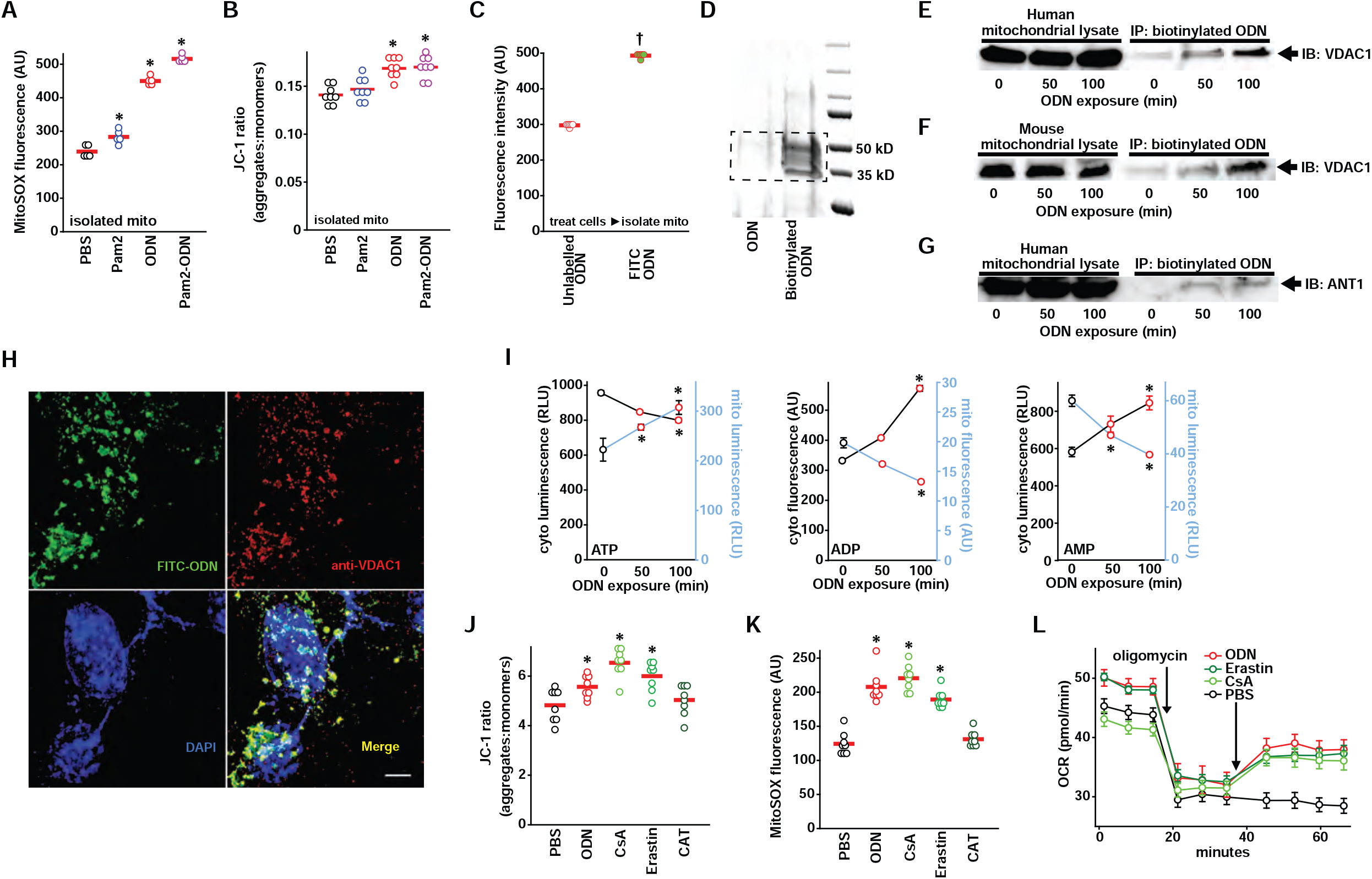
Blocking mitochondrial nucleotide transition by CpG ODN leads to generation of antimicrobial mtROS. (**A**) mtROS production and (**B**) Δ_Ψm_ increase in isolated mitochondria that were treated with Pam2, ODN or both. (**C**) Fluorescence intensity of mitochondria isolated from HBEC3-KT cells treated with FITC-labeled or unlabeled ODN. (**D**) Mitochondria were isolated from the biotinylated ODN-treated HBEC3-KT cells, and streptavidin precipitants from mitochondrial lysates were resolved by polyacrylamide gel electrophoresis and silver stained. Dashed line indicates bands excised for mass spectrometry analysis. As in **D**, streptavidin precipitants were probed for VDAC1 in mitochondrial lysates from (**E**) human or (**F**) mouse cells following treatment with ODN for the indicated time. Mitochondrial lysates from human cells were precipitated and probed for ANT1 (**G**) following treatment with ODN for the indicated time. (**H**) Representative images of HBEC3-KT cells treated with FITC-labeled ODN then stained with Alexa Fluor 555-labeled anti-VDAC1 antibody. Scale bar, 100 μm. (**I**) Measurements of cytosolic and mitochondrial levels of ATP, ADP & AMP in ODN-treated HBEC3-KT cells at the indicated times. Mitochondrial membrane potential Δ_Ψm_ (**J**) and mtROS (**K**) 100 min after HBEC3-KT treatment with the indicated mitochondrial permeability modulators. (**L**) Seahorse analysis of oxygen consumption following the indicated mitochondrial permeability modulators in oligomycin-inhibited HBEC3-KT cells, shown as mean ± SEM. * p<0.001 vs PBS by ANOVA; † p<0.001 vs unlabeled ODN by two-way Student’s t test. Mito, mitochondria; VDAC1, voltage dependent anion channel 1; ANT1, adenine nucleotide translocator 1; CsA, cyclosporin A; CAT, carboxyatractyloside; OCR, oxygen consumption rate.

To identify potential mitochondrial ODN binding partners, HBEC3-KT cells were treated with biotin-labeled ODN, then the streptavidin precipitants from mitochondria lysates were resolved on an SDS PAGE gel. The silver stain in Figure 3D demonstrates bands present only after ODN treatment. The boxed area was excised and liquid chromatography–mass spectrometry proteomic analysis generated a list of candidate targets.

Among the most differentially detected peptides was voltage-dependent anion-sensitive channel 1 (VDAC1), an outer mitochondrial membrane protein component of the VDAC1-ANT1-mCK complex that regulates exchange of ATP and ADP between the mitochondria and cytosol (25, 26). In targeted pulldown studies in mouse and human cells, we confirmed that VDAC1 was detected in immunoprecipitated samples of biotinylated ODN treated cells (Figure 3E-F, Figure S6). The association of ODN with VDAC1 increased with treatment time. We also detected association of ODN with adenine nucleotide translocator 1 (ANT1), the inner mitochondrial membrane component of the VDAC1-ANT1-mCK complex (Figure 3G, Figure S6). When HBEC3-KT cells were treated with fluorescently labeled ODN then VDAC1 was localized with fluorescently labeled antibody (Figure 3H), 75% of pixels occupied by VDAC1 were also occupied by ODN. The mean Pearson’s correlation coefficient between ODN and VDAC1 pixel intensity was 0.84 (Figure S7), indicating a high degree of colocalization in intact cells.

Given the role of the VDAC1-ANT1-mCK complex as an ATP:ADP antiporter, we investigated whether altered ATP/ADP localization might account for the changes in whole-cell energy stores previously observed following ODN treatment (Figure 2C-E). Indeed, we found that ODN treatment causes mitochondrial ATP levels to rapidly increase and cytosolic ATP levels to precipitously decline, with the opposite pattern for ADP and AMP (Figure 3I), consistent with a ODN-induced blockade of ATP:ADP antiporter function. To test whether VDAC1 antagonism can explain the ODN effect on mitochondrial energy metabolism, we investigated the effects of a known VDAC1 inhibitor erastin (27, 28) and an ANT1 inhibitor carboxyatractyloside (CAT) (29, 30). Because VDAC1 is one of the subunits composed of the mitochondrial permeability transition pore (mPTP), we also exposed cells to mPTP inhibitor cyclosporin A (31, 32). As shown in Figure 5J-L, erastin and cyclosporin A caused changes in Δ_Ψm_, mtROS production, and oxygen consumption that were comparable to ODN, suggesting that blocking VDAC-mediated mitochondrial nucleotide exerts these effects.

### AMPK-directed metabolic reprogram increases electron delivery to complex II

AMP-activated kinase (AMPK) is a cellular energy sensor that is activated when cytosolic AMP levels rise due to ATP consumption (33, 34), in turn activating catabolic pathways to promote ATP production, including acetyl-CoA carboxylase (ACC) (35, 36). Congurent with the observations that ODN alters cellular ATP/ADP/AMP stores (Figures 2-3), reverse phase protein array analysis of HBEC3-KT cells treated with ODN revealed AMPK and ACC to be among the most activated signaling pathways following ODN exposure (Figure 4A). We confirmed time-dependent ODN-induced phosphorylation of AMPKα1, AMPKα2 and ACC in vitro (Figure 4B, Figure S8). Similarly, AMPKα1 phosphorylation was induced by both erastin and cyclosporin A in vitro (Figure 4C, Figure S8). ODN-induced AMPKα phosphorylation was also demonstrated by immunofluorescence staining in mouse airways (Figure 4D-E). When AMPKα genes were conditionally deleted (Figure S9), ODN-inducible mtROS production was significantly reduced without impacting baseline mtROS production (Figure 4F).

**Fig. 4.**
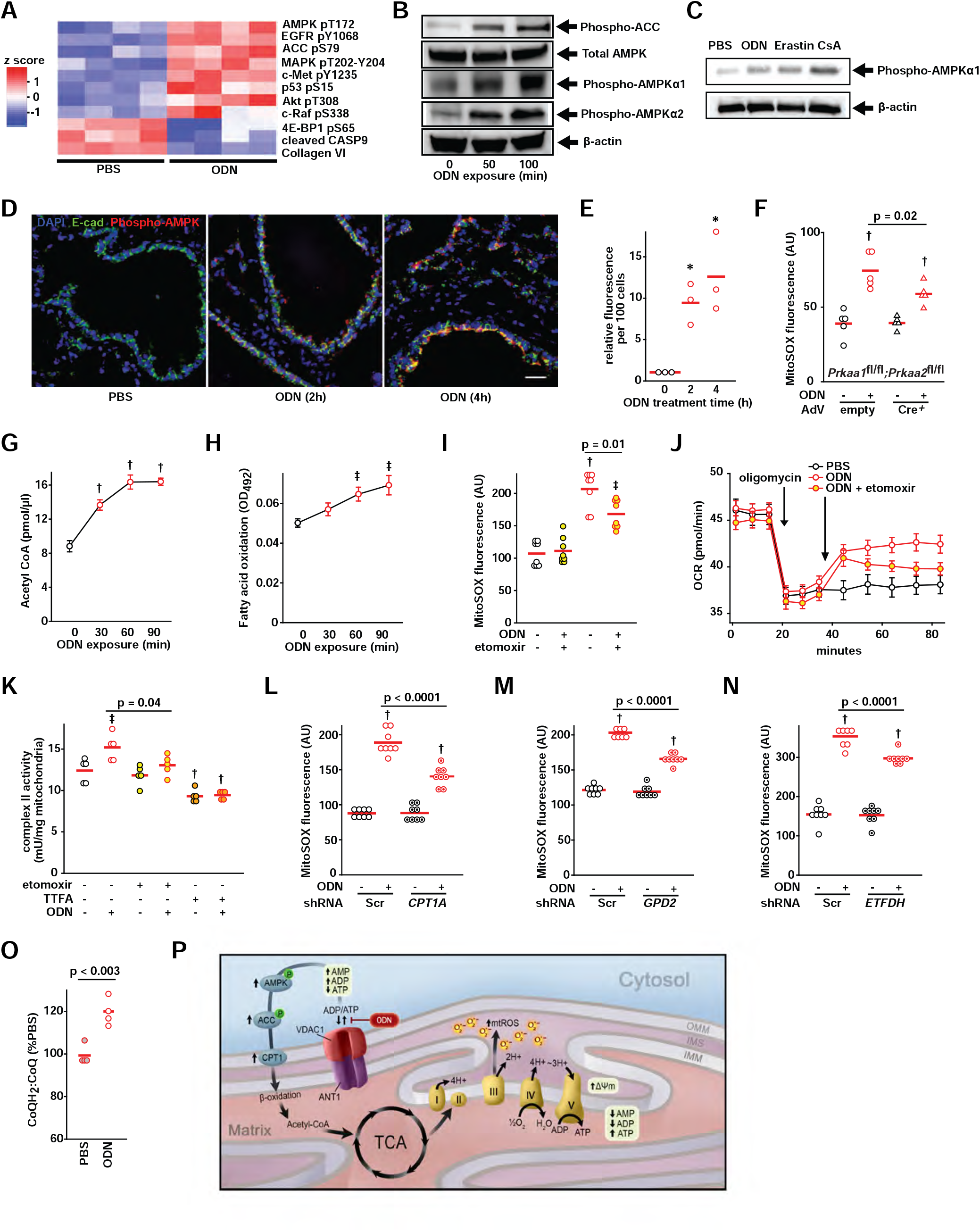
AMPK-regulated metabolic reprogram increases electron delivery to complex II. (**A**) RPPA heatmap from HBEC3-KT cells treated with PBS or ODN. (**B**) Immunoblot of AMPK and ACC proteins after ODN treatment. (**C**) Immunoblot for phospho-AMPKα1 following treatment with the indicated mitochondrial permeability modulators in HBEC3-KT cells. (**D**) Phospho-AMPK immunofluorescence in mouse lungs after treatment with ODN. Scale bar, 50 μm. (**E**) Quantification of fluorescence in **D**. (**F**) mtROS production in primary *Prkaa1*^*fl/fl*^*;Prkka2*^*fl/fl*^ mouse tracheal epithelial cells infected with empty or Cre^+^ adenovirus, then treated with PBS or ODN. (**G**) Acetyl-CoA levels in ODN-treated HBEC3-KT cells. (**H**) Fatty acid oxidation after ODN treatment. (**I**) mtROS production following treatment with ODN and/or β-oxidation inhibitor etomoxir. (**J**) Oxygen consumption following the indicated treatments, shown as mean ± SEM. (**K**) HBEC3-KT cell complex II activity following treatment with the indicated agents.ODN-induced mtROS production in cells with knockdowns of gene CPT1A (**L**) and the genes for electron shuttles GPD2 (**M**) or ETFDH (**N**). (**O**) Ratio of reduced:oxidized CoQ in mitochondria isolated from HBEC3-KT cells treated with PBS or ODN. (**P**) Schematic model of mtROS formation induced by ODN via metabolic reprogramming. * p <0.01 vs 0 min; † p <0.001 vs. (syngeneic) PBS treated; ǂ p < 0.02 vs (syngeneic) PBS treated. RPPA, reverse phase protein array; AMPK, AMP-activating protein kinase; ACC, acetyl-CoA carboxylase; AdV, adenovirus; OCR, oxygen consumption rate; Scr, scrambled shRNA control; CPT1A, carnitine palmitoyltransferase 1A; GPD2, glycerol-3-phosphate dehydrogenase 2; ETFDH, electron transfer flavoprotein-ubiquinone dehydrogenase.

A principle means by which AMPK-ACC pathway activation promotes mitochondrial energy production is through increased carnitine palmitoyltransferase 1 (CPT1)-dependent fatty acid β-oxidation. We found that ODN treatment increases acetyl CoA concentrations and fatty acid β-oxidation (Figure 4G-H). Further, treatment of HBEC3-KT cells with etomoxir, an irreversible inhibitor of CPT1 (37, 38), significantly attenuated ODN-induced mtROS production, oxygen consumption and ETC complex II activity (Figure 4I-K). Knockdown of *CPT1* (Figure S10) also attenuated inducible mtROS production to a similar degree to etomoxir (Figure 4L).

Mitochondrial fatty acid β-oxidation generates NADH and FADH_2_, which contribute electrons to the ETC via electron shuttle proteins. Specifically, FADH_2_-carried electrons are transferred to coenzyme Q (CoQ) by electron flavoprotein dehydrogenase (ETFDH) (39), while NADH-carried electrons are transferred to complex II by glycerol-3-phosphate dehydrogenase (GPD2) (40). Knocking down either of these shuttles (Figure S10) attenuated ODN-induced mtROS production (Figure 4M-N). Fatty acid β-oxidation also generates acetyl-CoA which transfers electrons to the ETC via the tricarboxylic acid (TCA) cycle. Treatments with TCA intermediate metabolites oxaloacetate and α-ketoglutarate or the analogue dimethyl malonate attenuated ODN-induced mtROS production (Figure S11). Dimethyl malonate and oxaloacetate are ETC complex II inhibitors while α-ketoglutarate inhibits glutaminolysis (41-43).

In the ETC, CoQH_2_ is generated when CoQ accepts electrons from FADH_2_. The CoQH_2_:CoQ ratio has been described as an indicator of ETC efficiency, with an increased ratio associated with increased mtROS production (44). As in Figure 4O, ODN treatment caused an increase in CoQH_2_:CoQ ratio. Although augmented β-oxidation is required for maximal ODN-induced mtROS production, inducible mtROS production can also be partially attenuated by inhibiting glycolysis and/or glutaminolysis (Figure S12). Future work will explore the contributions of these pathways.

Together, these results indicated that ODN-induced and AMPK-regulated metabolic reprogramming enhances electron delivery to ETC, increases complex II activity and eventually drives mtROS induction. These findings were schematically summarized in Figure 4P.

### ODN-induced mtROS generation at complex III is ΔΨm-dependent

It is intriguing that ODN simultaneously increases complex II activity and decreases complex III activity. We hypothesized that these changes in ETC complex activity might provide insights into the site(s) of mtROS generation. A series of ETC complex inhibitors were used to determine the roles of ETC complexes in ODN-induced mtROS generation.

mtROS are formed when molecular oxygen interacts with an electron leaked among electron transport chain (23), typically from complex I [flavin (F) site and ubiquinone reduction (Q) site] or complex III (Q_o_ site) (45). Whereas mtROS formation at complex I following other stimuli can be inhibited by rotenone (Q site) or diphenyleneiodonium (F site) (46, 47), neither agent impeded ODN-induced mtROS (Figure S13). Similarly, while TCA cycle input can support reverse election transport from complex II to complex I in the setting of high Δ_Ψm_ (48), we found that neither succinate nor fumarate influence ODN-induced mtROS formation (Figure S11). These findings, along with the modest ODN impact on complex I activity (Figure 2A, Figure S2), indicated that complex I is not a major site of ODN-induced mtROS production. In contrast, while inhibitors of complex II activity reduced ODN-induced mtROS, inhibitors of complex III enhanced ODN-induced mtROS dramatically (Figure S13). We thus concluded that complex III is the main ODN-induced mtROS generating site following forward electron transfer.

Additionally, while complex IV inhibition had no impact on mtROS induction, the complex V inhibitor oligomycin decreased ODN-induced mtROS production. As oligomycin treatment collapses mitochondrial membrane potential Δ_Ψm_, this result suggested that ODN-induced Δ_Ψm_ increases may be required for mtROS formation.

We compared the impacts of ODN and complex III inhibitor antimycin on complex III activity and mtROS induction. Although treatment with antimycin or ODN resulted in similar inhibition of complex III enzymatic activity and initial mtROS production (Figure 5A-B), the two agents function differently. When stigmatellin and myxothiazol inhibit electron transfer from complex III to complex IV by binding the CoQH_2_ (ubiquinol) oxidation (Q_o_) site, antimycin cannot induce further oxygen consumption for superoxide production while ODN still can (Figure 5C). Antimycin can induce rapid induction of mtROS production in intact epithelial cells and isolated mitochondria, but this effect plateaus by 40 min. Conversely, ODN-induced mtROS continued to increase throughout the period of exposure in intact cells and isolated mitochondria, suggesting different mechanisms of mtROS generation (Figure 5D-E). Central to these differences appeared to be their opposing effects on Δ_Ψm_. In both whole cell and isolated mitochondria models, ODN induced increased Δ_Ψm_ while antimycin reduced Δ_Ψm_ below that observed in sham-treated samples (Figure 5F-G). Disrupting Δ_Ψm_ with the uncoupler FCCP (Figure 5H-I) significantly impaired ODN-induced mtROS generation and but had little effect on antimycin-induced mtROS generation (Figure 5J-K). FCCP treatment demonstrated that ODN-induced oxidation of cytochrome b_H_ is Δ_Ψm_-dependent, whereas FCCP did not alter the oxidation of cytochrome b_H_ in antimycin-treated mitochondria (Figure 5L). Congruently, in isolated mitochondria, FCCP reversed ODN-impaired complex III electron transfer activity but had no such effect on antimycin treated mitochondria (Figure 5M). The generation of ODN-induced mtROS at complex III is graphically displayed in Figure 5N. Under homeostatic conditions, complex III quickly transfers CoQH_2_-carried electrons to cytochrome c1, which, in turn, transfers the electrons to complex IV. This process facilitates proton pumping across the inner mitochondrial membrane and establishes normal Δ_Ψm_ (49, 50). However, ODN-induced increase in Δ_Ψm_ hinders the proton pumping, impeding electron transfer at the Q_o_ and quinone reduction (Q_i_) sites. This Δ_Ψm_-dependent retardation of electron transfer in complex III, in coordination with an increased forward electron transfer from complex II, increases the likelihood that highly-reactive electrons will “leak” to interact with free oxygen, resulting in increased formation of mtROS, in the form of superoxide, at complex III (51-53).

**Fig. 5.**
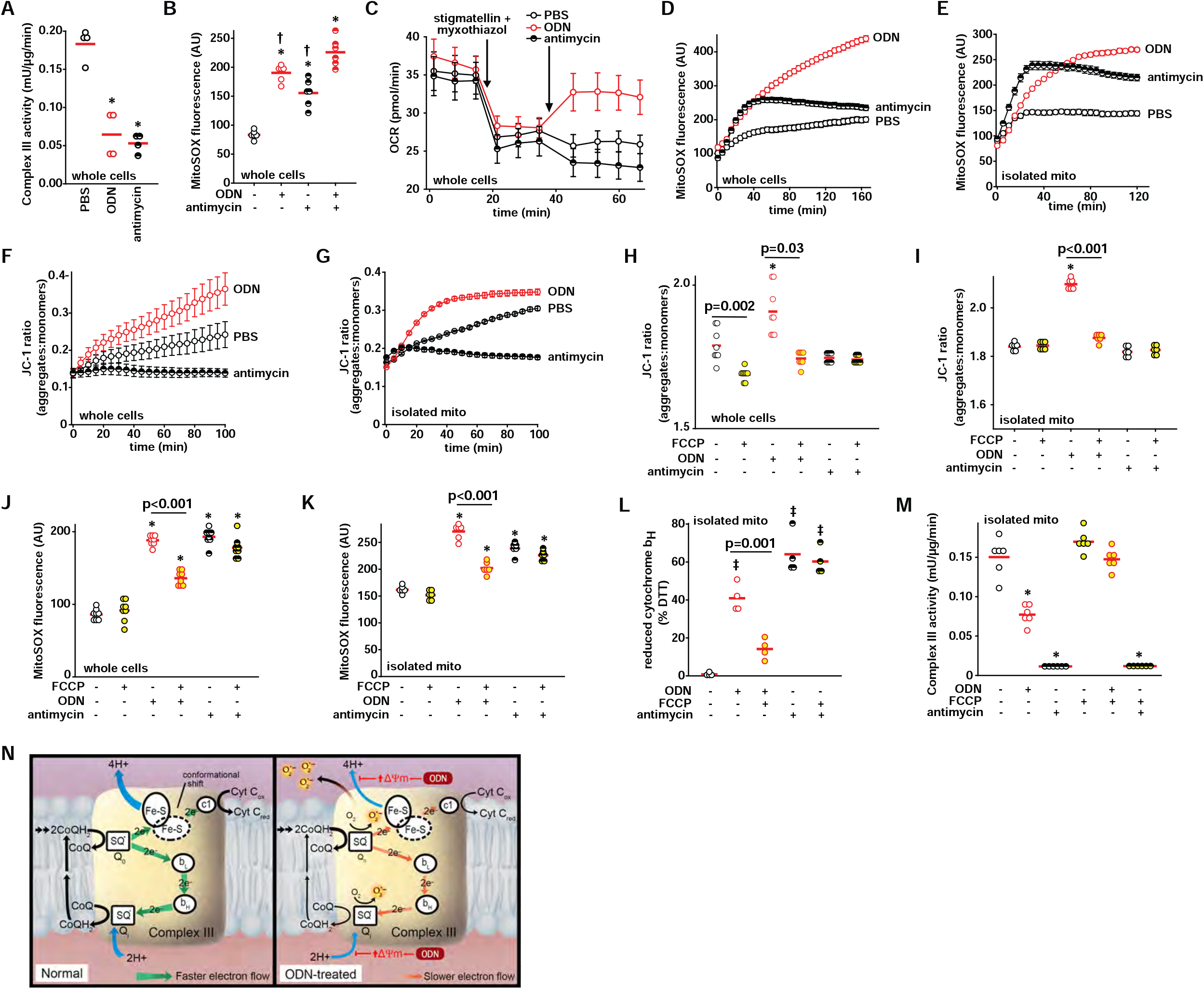
mtROS formation at complex III is ΔΨm-dependent. (**A**) Electron transport chain complex III activity in HBEC3-KT cells 100 min after ODN or antimycin treatment. (**B**) mtROS production 100 min after the indicated treatments. (**C**) Oxygen consumption following the indicated treatments in stigmatellin and myxothiazol-inhibited HBEC3-KT cells, shown as mean ± SEM. Time course of mtROS generation following PBS, ODN or antimycin treatment in (**D**) HBEC3-KT cells or (**E**) isolated mitochondria. Time-dependent mitochondrial membrane potential in (**F**) HBEC3-KT cells or (**G**) isolated mitochondria. Mitochondrial membrane potential Δ_Ψm_ 100 min after ODN or antimycin treatments in (**H**) HBEC3-KT cells or (**I**) isolated mitochondria with or without FCCP pre-treatment. mtROS generation 100 min after ODN or antimycin treatments in (**J**) HBEC3-KT cells or (**K**) isolated mitochondria with or without FCCP pre-treatment. (**L**) Reduced mitochondrial complex III cytochrome b_H_ levels following ODN or antimycin treatment with or without FCCP pre-treatment, expressed relative to DTT-treated mitochondria (DTT-treated presumed 100% reduced). (**M**) Complex III activity in isolated mitochondria 15 min after the indicated treatments. (**N**) Schematic model of Δ_Ψm_-dependent mtROS formation at complex III. * p<0.001 vs PBS by ANOVA; † p<0.008 vs ODN + antimycin treated by ANOVA; ǂ p<0.02 vs PBS by ANOVA. mito, mitochondria; DTT, dithiothreitol.

Thus, while dissecting the process of ODN-induced antimicrobial mtROS formation, we identified that mtROS induction requires both AMPK-directed metabolic reprograming to augment electron delivery to ETC complex II (Figure 4P) and increased Δ_Ψm_ to retard electron transfer at complex III (Figure 5N).

### mtROS induction stimulates TLR9-independent antimicrobial effects

To demonstrate the protective effects of antimicrobial mtROS induced by ODN, we have shown that scavenging mtROS by mitoTEMPO or mitoQ (Figure S14) significantly decreases the bacterial killing induced by Pam2-ODN combined treatment in HBEC3-KT cells (18). While pre-treatment with either an ETC complex II inhibitor TTFA or the Δ_Ψm_ uncoupler FCCP alone inhibited ODN-induced mtROS to some extent, TTFA-FCCP combination treatment maximally inhibited mtROS production in HBEC3-KT cells (Figure S15) and reversed the mitochondrial reduced:oxidized ratio of ODN-treated cells (Figure 6A-B). As shown in Figure 6C and Figure S15, the bacterial killing induced by Pam2 and ODN was obviated in HBEC3-KT cells when the cells were pretreated with TTFA-FCCP. Congruently, wild type mice with impaired lung epithelial mtROS generation due to aerosolized TTFA-FCCP pretreatment prior to treatment with Pam2 and ODN were less protected against *P. aeruginosa* pneumonia than were mice who received sham aerosol pretreatment prior to receiving Pam2-ODN (Figure 6D). Notably, both the Pam2-ODN-induced *P. aeruginosa* pneumonia protection and the TTFA-FCCP-induced impairment were observed in *Tlr9* knockout mice (Figure 6E). Consistent with our prior reports (17), airway delivery of TTFA-FCCP had no observed systemic effects on the mice in the absence of infection (Figure 6E). In light of our data supporting VDAC1 blockade as central mediator of protection, we tested whether erastin substituted for ODN could protect against infection. In combination with Pam2, erastin and ODN induced comparable bacterial killing by HEBC3-KT cells (Figure 6F). Strikingly, when delivered by aerosol with Pam2, erastin yielded a similar survival advantage to ODN following *P. aeruginosa* pneumonia challenge in mice (Figure 6G).

**Fig. 6.**
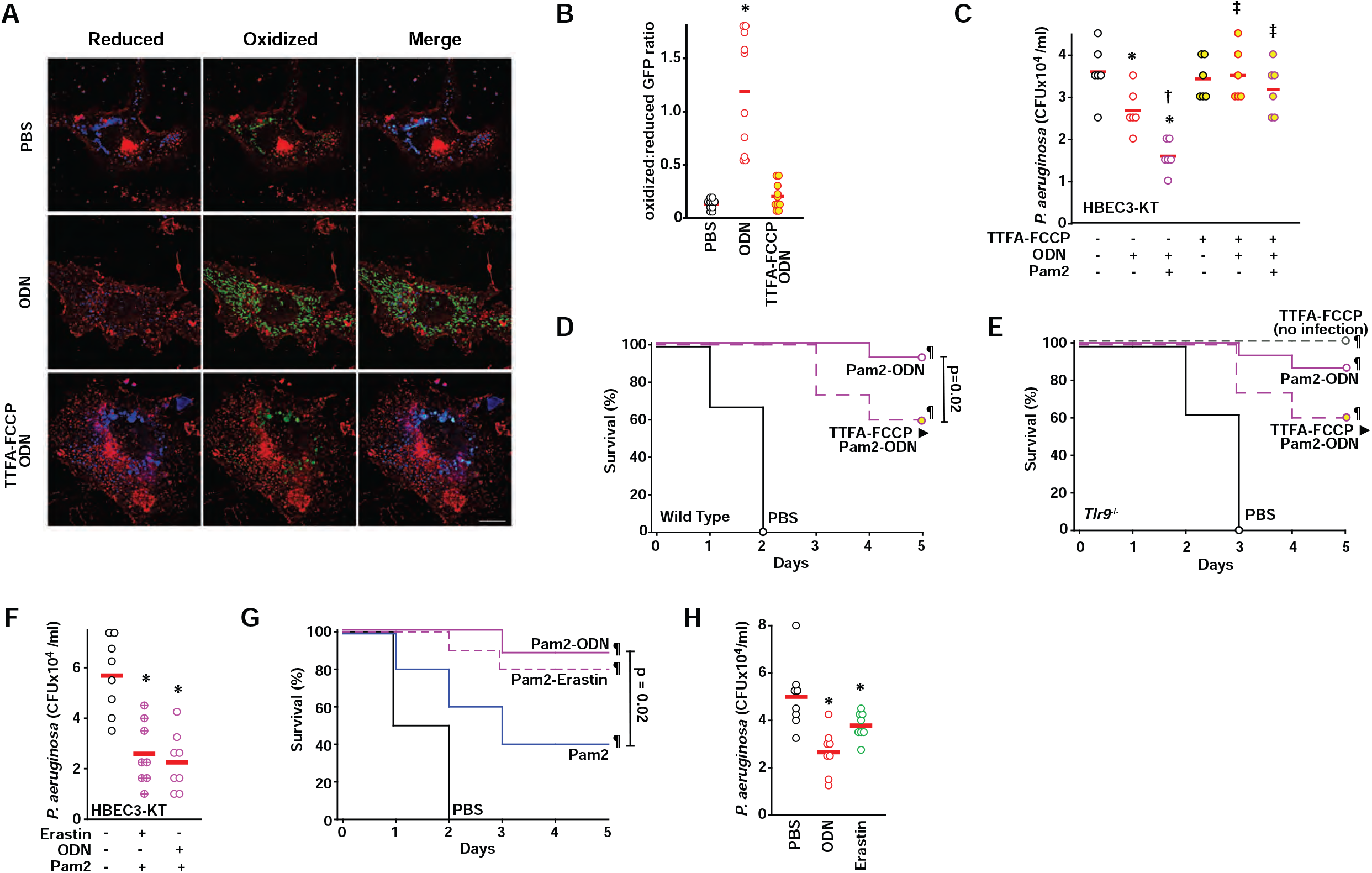
mtROS induction stimulates antimicrobial responses. (**A**) Representative fluorescence images primary tracheal epithelial cells harvested from mt-roGFP mice, pre-treated (or not) with TTFA and FCCP, then treated with PBS or ODN. Images shown as gradient of color intensity from the reduced (blue) form to the oxidized (green) form of roGFP. Scale bar, 50 μm. (**B**) Ratio of the fluorescence intensity of the oxidized:reduced roGFP from **A**, quantified at 488 nm and 405 nm, respectively. (**C**) Bacterial burden of HBEC3-KT cells treated with the indicated ligands with or without TTFA-FCCP treatment. (**D**) Survival of wild type mice challenged with *P. aeruginosa* one day after nebulized treatment with PBS or Pam2 and ODN with or without TTFA-FCCP (n=15 mice/group). (**E**) Survival of *Tlr9*^-/-^ mice challenged with *P. aeruginosa* one day after nebulized treatment with PBS or Pam2 and ODN with or without TTFA-FCCP (n=15 mice/group). (**F**) Bacterial burden of HBEC3-KT cells treated with Pam2 and erastin or ODN. (**G**) Mouse survival of *P. aeruginosa* challenge given one day after nebulized treatment with the indicated agents (n=15 mice/group). (**H**) Mouse lung bacterial burden immediately after *P. aeruginosa* challenge following treatment with the indicated agents (n=4 mice/group). * p <0.02 vs PBS, † p< 0.05 vs ODN, ‡ p < 0.05 vs same ligand without TTFA-FCCP, ¶ P <0.0001 vs. PBS.

Even when delivered without Pam2, both ODN and erastin induced significant reductions in the lung bacterial burden (Figure 6H).

In summary, we found that antimicrobial mtROS are generated from lung epithelial cells following ODN treatment. ODN interacts with VDAC1 to alter mitochondrial nucleotide transport, driving AMPK-ACC-CPT1-mediated electron delivery to ETC complex II and increasing Δ_Ψm_ to promote superoxide production at complex III.

## DISCUSSION

Synthetic CpG ODNs have been explicitly developed as immunomodulators and adjuvants (24, 54, 55). ODN CpG motifs mimic naturally-occurring pathogen-associated molecular patterns recognized by TLR9 that initiate NF-κB-dependent antimicrobial signaling cascades (56-58). Here, we report previously unknown, TLR9-independent induction of immunometabolic reprogramming by ODN that results in generation of pneumonia-protective mtROS.

ODN-induced mtROS formation is fundamentally a manifestation of altered energy metabolism. ODN interacts with VDAC1 and ANT1 localized in the mitochondrial membranes. The VDAC1-ANT1-mCK complex regulates the exchange of metabolites between the mitochondria and cytosol (25, 26). We show that VDAC1 binding ODN perturbs cellular nucleotide distribution, activating the AMPK-ACC pathway, and promoting fatty acid β-oxidation. Fatty acid β-oxidation augments FADH_2_-carried electron flux to the ETC by ETFDH, GPD2, and the TCA cycle, all converging on the CoQ pool (CoQH_2_ and CoQ). Disruption of any of these elements leads to decreases in inducible mtROS generation.

The increased electron delivery as a consequence of AMPK-ACC activation results in increased complex II activity, however, these observations do not resolve why complex III activity are decreased. During normal bifurcated electron transfer in complex III (59, 60), semiquinone (SQ•-) intermediates forming at the Qo or Qi site instantaneously transfer electrons to the low potential heme b (bL) or CoQ, minimizing electron leakage. However, under certain conditions, accumulation of SQ•^-^ increases electron leak (51). In one example, antimycin inhibits CoQ reduction at the Q_i_ site, leading to electron accumulation on cytochrome b hemes, allowing SQ•^-^ more time to interact with molecular oxygen to form superoxide at the Q_o_ site (61, 62).

Alternatively, high Δ_Ψm_ attenuates the proton pump, retarding electron transfer and sustaining cytochrome b hemes in reduced states that cause accumulation of SQ•^-^ at the Q_o_ and Q_i_ sites (52, 63). Here, our findings support the latter as the responsible mechanism as uncoupling Δ_Ψm_ with FCCP reduces ODN-induced mtROS and reverses ODN-impaired complex III activity.

Thus, Δ_Ψm_ accentuation by VDAC-perturbed ATP accumulation in mitochondria increases mtROS production and impairs complex III activity.

Although ROS production is often regarded as an untoward cellular event that contributes to degenerative diseases (64, 65), there is robust evidence that controlled mtROS generation contributes to critical signaling events in a wide range of physiologic processes that extend host survival (66-70), including by augmentation of protective antimicrobial responses (71-73). Superoxide formed at the complex III Q_o_ site may be particularly well suited to function as a cytosolic signaling molecule, as the Q_o_ site is adjacent to the intermembrane space with about half of its superoxide diffusing to the cytoplasmic side of the inner membrane (74, 75).

Here, we demonstrate that complex III-dependent mtROS induction is required for maximally ODN-induced bacterial killing in vitro and in vivo. Although the current work does not explicitly test whether mtROS directly kill pathogens or act as cell signals to initiate antimicrobial responses, both topics are areas of active investigation now.

In summary, we identify metabolic mechanisms underlying the ODN-induced antimicrobial mtROS formation. Under physiologic conditions, mtROS production is exquisitely tightly regulated, but we show here that therapeutic manipulation of mtROS is achievable, protective against otherwise lethal infections, and well tolerated by the host. Indeed, this intervention has also been safely tested in five completed human trials (NCT04313023, NCT04312997, NCT03794557, NCT02566252, NCT02124278) with more in preparation.

Because of our interest in pneumonia, all of the current work is performed in lung epithelial cells, but we anticipate similar responses can be detected in other epithelial cells and, likely, other cell types.

## METHODS

### Primary Cell Cultures and Cell lines

To isolate mouse tracheal epithelial cells (mTECs), mice were anesthetized and tracheas were excised and digested in 1.5 mg/ml Pronase overnight at 4 ºC. mTECs were harvested by centrifugation and then cultured on collagen coated tissue culture plates or transwells in Ham’s F12 media supplemented with differentiation growth factors and hormones as previously described (76).

Normal human bronchial epithelial (NHBE) cells were purchased from American Type Culture Collection (ATCC, Manassas, VA) and cultured in airway epithelial cell basal medium supplemented with bronchial epithelial cell growth kit (ATCC, Manassas, VA).

Immortalized Human bronchial epithelial (HBEC3-KT) cells were kindly provided by Dr. John Minna. Murine lung epithelial (MLE-15) cells were kindly provided by Dr. Jeffrey Whitsett.

HBEC3-KT and MLE-15 cells were authenticated by the UT MD Anderson Characterized Cell Line Core Facility and IDEXX Bioresearch (Columbia, MO), respectively. HBEC3-KT cells were cultured in keratinocyte serum-free medium supplemented with human epidermal growth factor and bovine pituitary extract (Thermo Fisher Scientific, Grand Island, NY). MLE-15 cells were cultured in DMEM/F2 medium supplemented with 2% of fetal bovine serum and 0.5% of Insulin-Transferrin-Selenium (Thermo Fisher Scientific, Grand Island, NY). Cell cultures were maintained in the presence of 1% of penicillin/streptomycin and glutamine. All cells were cultured at 37 °C with 5% CO_2_. All human cell experiments were performed in accordance with Institutional Review Board of The University of Texas MD Anderson Cancer Center (MDACC).

### Mice

Wild type C57BL/6J mice were purchased from The Jackson Laboratory (Bar Harbor, ME). *Prkaa1*^*fl*^ and *Prkaa2*^*fl*^ mice were purchased from Jackson. *TLR9*^***–/–***^ mice were provided by Dr. Shizuo Akira (77). *CMV mt-roGFP* mice were generated by Dr. D James Surmeier and kindly provided by Dr. Farhad Danesh (19, 20). *Sftpc-Cre* mice were kindly provided by Dr. Brigid Hogan (78). All mouse experiments were performed in accordance with the MDACC Institutional Animal Care and Use Committee.

### In Vivo Infection Model

As previously described (8, 18), 10 ml of combined 4 µM Pam2CSK4 and 1 µM ODN M362 in 1× phosphate-buffered saline (PBS) was placed in an Aerotech II nebulizer (Biodex, Shirley, NY) and delivered to unrestrained mice in an exposure chamber via an influx polyethylene tube. Nebulization was driven by 10 L/min air supplemented with 5% CO_2_. The exposure chamber connects with an identical efflux polyethylene tube with a low resistance microbial filter (BB50T, Pall, East Hills, NY) at its end vented to a biosafety hood.

*Pseudomonas aeruginosa* strain PA103 was purchased from ATCC and stored as frozen stock (1×10^8^ colony-forming unit CFU/ml) in 20% glycerol in Luria-Bertani (LB) medium. Typically, 1 ml of frozen stock was incubated overnight in 100 ml of Tryptic Soy Broth (TSB) at 37°C with 5% CO_2_, then expanded in 1 liter of fresh LB media at 37°C to OD 600 of 0.52. Bacterial suspensions were centrifuged, washed, re-suspended in 1× PBS, and aerosolized using the same nebulization system for Pam2-ODN treatment. For all bacterial challenges, a nebulized inoculum of 10 ml of ∼2×10^10^ CFU/ml were delivered. If not specified, 6 to 8 weeks old single sex mice were used for in vivo infection conducted in a BSL2 biohazard lab. Immediately following bacterial challenge, some mice were anesthetized and mouse lungs were harvested and homogenized using a Mini-Beadbeater-1 (Biospec, Bartlesville, OK). The lung homogenates were used to count lung colony-forming units (CFUs). The remaining mice were closely monitored for 12 days. The relevant euthanasia-triggering criteria consist of any evidence of distressed behaviors including hypothermia, impaired mobility, respiratory distress, and inability to access food or water. When mice were identified to meet the criteria, they were subjected to euthanasia immediately. At least 8-10 mice per condition were evaluated for survival analysis, 4 mice per condition were sacrificed for pathogen burden assessment. Challenges were performed a minimum of 3 times.

### In Vitro Pathogen Killing Assay

HBEC3-KT cells or MLE-15 cells were cultured on 6-well plates in complete media until cell growth reached ∼80% confluence. Cells were replaced with fresh, antibiotic-free media containing PBS, Pam2, ODN or Pam2-ODN. The final concentrations of Pam2 or ODN in media were 2.4 µM or 0.6 µM, respectively. 4 h after the treatment, 20 µl of *P. aeruginosa* PA103 (1×10^5^ CFUs/ml) were added to each culture well. 4 h after bacteria inoculation, 20 µl of supernatant from each well was aspirated, serially diluted, plated on a TSB agar plate and incubated for 16 h at 37 °C. Bacterial CFUs were counted after the incubation (18). Studies were performed a minimum of 3 times with 4 biological replicates per condition.

### Mitochondrial ROS Detection, Scavenging and Inhibition

To detect mtROS generation, cells were incubated with 5 µM of each indicated detector, MitoSOX red, ROSstar 550, MitoTracker Red CMXRos or tetramethylrhodamine (TMRM) in a black-walled, clear bottomed 96-well plate for 1/2 h before ODN or PBS treatment (17). After fluorescent mtROS detectors were washed off, fluorescence was continuously measured on a BioTek Synergy2 plate reader for 3 h immediately after ODN or PBS addition.

Excitation/emission wavelengths for mtROS-detecting agents are 510 nm/580 nm. Studies were performed a minimum of 3 times with 8 biological replicates per condition.

To scavenge mtROS, HBEC3-KT cells were exposed to 100 nM MitoTEMPO or 10 µM MitoQ for 1 h prior to fluorescent mtROS detector incubation and ODN or PBS treatment.

To disrupt mtROS production, HBEC3-KT cells were exposed to compounds that inhibit mitochondria electron transport chain activity. These include rotenone (10 μM), TTFA (200 μM), atpenin (10 nM), antimycin (100 nM & 5 μM), sodium azide (1 μM), oligomycin (2 μM) and FCCP (400 nM) etc., for 1 h prior to fluorescent mtROS detector incubation and ODN or PBS treatment. Inhibition of ODN-induced mtROS generation in vitro was achieved by concurrent application of TTFA (200 μM) and FCCP (200 nM).

To inhibit mtROS generation in vivo, mice were exposed to TTFA (200 mM) and FCCP (800 µM) in 10 ml of 50% DMSO solution in 1× PBS by nebulization. The 50% DMSO solution was nebulized as vehicle control. Mice received TTFA-FCCP or vehicle control were treated with either Pam2-ODN or PBS and then subjected to *P. aeruginosa* PA103 challenge.

### JC-1 Assay

To monitor changes in mitochondria membrane potential (Δψ_m_) upon ODN treatment, HBEC3-KT cells were incubated with 2 µM of JC-1 fluorescence dye for 30 min. After the JC-1 dye was washed off, fluorescence was continuously measured with ODN or PBS treatment using a BioTek Synergy2 plate reader at wavelengths of 510nm/580nm for J-aggregates and 490nm/525nm for J-monomers. A higher ratio of J-aggregates to J-monomers indicates a higher Δψ_M_. To inhibit mitochondrial membrane polarization, TTFA (200 μM) and FCCP (200 nM) were concurrently applied for 1 h before JC-1 incubation and ODN or PBS treatment. Studies were performed a minimum of 3 times with 8 biological replicates per condition.

### Indirect Immunofluorescence Assay and Co-localization Image Analysis

PBS or ODN-treated HBEC3-KT cells or MLE-15 cells growing on a chambered coverglass or frozen lung sections from PBS or ODN-treated mice were fixed with 2% paraformaldehyde, permeabilized with 0.1% Triton X-100, and blocked with 2% goat serum in 1× PBS. Cells or lung sections were incubated with primary antibodies against Phospho-AMPKα-1,2 or VDAC1 at a dilution of 1:200 for 1 h, then with AlexaFluor secondary antibodies (Life Technologies, Carlsbad, CA) at a dilution of 1:500 for half an hour, and counterstained with 4′,6-diamidino-2-phenylindole (DAPI) for 15 minutes. Cells were visualized using a DeltaVision deconvolution fluorescence microscope (GE Life Sciences). Fluorescence intensity of microscope images was quantified using ImageJ.

Pixel intensity data of VDAC1 and ODN-FITC in the fluorescence images were imported into MATLAB and all analysis was performed using default settings. Confidence intervals were set at the 95% confidence limit. Pixel intensity values for VDAC1 and ODN-FITC were compared directly on a scatterplot. Six independent analyses were performed. A simple linear regression model using the least squares standard approach was fit to the data. Pearson’s correlation coefficients were calculated to determine whether VDAC1 pixel intensity tended to accumulate with ODN-FITC pixel intensity.

### Live Cell Fluorescence Imaging

Fluorescent measurement of intracellular mtROS generation was carried out on live and metabolically active mTECs generated from *CMV mt-roGFP* mice. Cells growing on collagen coated chambered coverglass were mounted onto microscopic chamber at 37 °C in air with 5% CO_2_, treated with ODN or PBS and washed with 1× live cell imaging buffer (Life Technologies). CellMask DeepRed was added to stain the cell membrane. Images were obtained using a DeltaVision deconvolution fluorescence microscope (GE Life Sciences) at excitation wavelengths of 405nm and 488nm at 100 minutes post ODN treatment. Fluorescence intensity of microscope images was quantified using ImageJ software.

### Measurement of Oxygen Consumption Rate

Oxygen consumption rate (OCR) was measured using a Seahorse XFe96 extracellular flux analyzer (Agilent, Santa Clara, CA). HBEC3-KT cells (1.5×10^4^ per well) were seeded into a XFe96 microplate and grew overnight in complete media. 1 h before the assay, the media was changed to Seahorse XF assay media (Agilent) and incubated in a non-CO_2_ incubator at 37 °C. The microplate was loaded onto the analyzer and basal respiration in these cells were recorded by real-time measurement of OCR. Then 25 μl of PBS, ODN and/or mitochondria inhibitors prepared in the assay medium were sequentially injected into each culture well (in 150 μl of assay media) via drug delivery ports. The final working concentrations of these testing regents in culture wells are ODN 1.2 µM, oligomycin 10 nM, and antimycin 1 µM. After injection of each regent, OCR was again measured. At the end of the assay, the number of viable cells was determined using trypan blue. OCR measurements were normalized to final cell numbers. OCR is expressed in pmole min^−1^. Studies were performed a minimum of 3 times with 12 biological replicates per condition.

### Western Blotting and Immunoprecipitation

HBEC3-KT cells or MLE-15 cells were suspended in NP-40 lysis buffer containing Halt protease and phosphatase inhibitor cocktail (Millipore), disrupted by sonication, and extracted at 4°C for 30 min. The protein concentration of the lysate was determined using bicinchoninic acid (BCA) protein assay (Pierce). 50 µg protein in 1× Laemmli buffer was separated by SDS-PAGE and then transferred onto polyvinylidene difluoride (PVDF) membranes (Millipore). The PVDF membranes were blotted with primary antibodies, detected by secondary antibodies with conjugated horseradish peroxidase, and developed using a Pico-sensitive chemiluminescence kit (Pierce). All membranes were stripped and re-blotted for β-actin or GAPDH as loading control.

Whole cell or mitochondria lysates were prepared with biotinylated ODN-treated HBEC3-KT cells or MLE-15 cells. To precipitate proteins Bound by biotinylated ODN in vivo, streptavidin beads (Pierce) were incubated with whole cell or mitochondria lysates containing 300 µg protein overnight at 4° C under constant gentle rotating. After incubation, streptavidin beads were centrifuged, washed with 1× PBS containing 0.05% Tween-20, resuspended in 50 μl of 2× SDS loading buffer, and boiled for 10 minutes. Elutes from the streptavidin beads were loaded onto SDS PAGE gel (Bio-Rad) and immunoblotted with VDAC1 or ANT1 antibody.

### Lentiviral shRNA Knockdown

GIPZ *E. coli* clones containing human *CPT1A, GPD2* and *ETFDH* lentiviral shRNA vectors were purchased from GE Dharmarcon (Lafayette, CO). The lentiviral shRNA vectors were purified using a QIAGEN plasmid kit. Lentiviruses bearing human *CPT1A, GPD2* or *ETFDH* shRNA were produced by co-transfection of the lentiviral shRNA vectors and lentiviral packaging vectors in 293T cells. The shRNA lentiviruses were collected and added into HBEC3-KT cell culture. Lentivirus-infected HBEC3-KT cells were selected by cell sorting based on GFP expression 3 days after infection. Efficiency of the shRNA knockdown was determined by immunoblot using anti-human *CPT1A, GPD2* or *ETFDH* antibodies.

### Adenoviral Cre Knockout

Adenovirus containing Cre recombinase or control empty vectors were purchased from Viral Vector Core at the University of Iowa. As previously described (79), adenoviral infections were carried out in mTECs derived from *Prkaa1*^*fl/fl*^*;Prkaa2*^*fl/fl*^ mice.

### Mitochondria Isolation

Mitochondria were isolated from either excised mouse lungs or harvested HBEC3-KT or MLE-15 cells. As previously described (80), a Polytron homogenizer (Pro Scientific, Oxford, CT) was used to dissociate tissues and cells. Mitochondria were separated from cytosol via serial centrifugation at 4 °C. The concentration of the isolated mitochondria was normalized in between PBS or ODN treatment groups using BCA protein assay (Pierce). The isolated mitochondria were maintained in mitochondrial isolation buffer on ice for further analysis.

### Biochemical Analysis of Mitochondrial Complex Activity

As previously described (81), spectrophotometric analysis of the respiratory chain complexes was performed on HBEC3-KT cells collected at the indicated time points after ODN treatment. The electron transport chain enzymes were assayed at 30° C using a temperature-controlled spectrophotometer (Ultraspec 6300 pro, Biochrom Ltd., Cambridge, England). The activities of complex I (NADH:ferricyanide reductase), complex II (succinate dehydrogenase), rotenone sensitive complex I+III (NADH:cytochrome c reductase), complex II + III (succinate:cytochrome c reductase) and complex IV (cytochrome c oxidase) were measured using appropriate electron acceptors/donors (82, 83). The increase or decrease in absorbance of cytochrome c at 550 nm was measured for complex I + III, II + III, or complex IV. The activity of NADH:ferricyanide reductase was measured by oxidation of NADH at 340 nm. For succinate dehydrogenase, the reduction of 2,6-dichloroindophenol (DCIP) was measured at 600 nm. For citrate synthase, the reduction of dithionitrobenzoic acid (DTNB) was measured at 412 nm. Enzyme activities are expressed in nmol/min/mg protein. Complex III activity was measured using a Mitochondrial Complex III activity assay kit (BioVision). Complex V activity was measured using a Quantichrom ATPase assay kit (Bioassay Systems).

### Luminescence Glo Assay

HBEC3-KT cells (1×10^4^ per well) were plated in an opaque-walled 96-well plate and grown overnight. Cells were treated with 0.6 µM ODN and collected at various time points after treatment. Cells were lysed on the plate and incubated with luminescence glo regents per the luminescence glo assay kit’s instructions (Promega). Luminescence was recorded using a BioTek Synergy2 plate reader.

### Fatty Acid Oxidation Assay

A non-radioactive fatty acid oxidation (FAO) assay kit (Biomedical Research Service, State University of New York at Buffalo) was used to measure FAO activity in ODN-treated HBEC3-KT cells. Cells were lysed and incubated with substrate octanoyl-CoA and then FAO assay solution in a 96-well plate, following the manufacturer’s instruction. Oxidation of the octanoyl-CoA by the cell lysate generates NADH, which is coupled to the reduction of the tetrazolium salt INT to formazan in the FAO assay solution. The absorbance of the newly formed formazan (at 490 nm) is proportional to FAO activity.

### Coenzyme Q_10_ Measurement by Triple Quadruple LC-MS

Levels of ubiquinone (CoQ_10_) and ubiquinol (reduced form) in PBS or ODN treated HBEC3-KT cells were measured using an Agilent 6460 triple quadruple mass spectrometer coupled with an Agilent 1290 series HPLC system. PBS or ODN treated cells were quickly washed with ice-cold PBS and then liquid nitrogen was poured onto cells to rapidly quench metabolic and chemical reactions. To extract ubiquinone and ubiquinol, 100 µl of 100% isopropanol were added and mixed with the cells. Coenzyme Q_9_ was added as an internal standard. The cell extracts were vortexed, centrifuged at 17,000 g for 5 minutes at 4 °C, and supernatants were transferred to clean auto sampler vials for direct injection. The mobile phase is methanol containing 5 mM ammonium formate. Ubiquinone, ubiquinol and CoQ_9_ were separated on a Kinetex® 2.6 µm C18 100 Å, 100 × 4.6 mm column. The flow rate was 700 µL/min at 37 °C. The mass spectrometer was operated in the MRM positive ion electrospray mode with the following transitions: Ubiquinone/oxidized, m/z 880.7→197.1; Uniquinol/reduced, m/z 882.7→197.1; CoQ_9_ (IS), m/z 795.6 →197.1. Raw files were imported and analyzed using Agilent Mass Hunter Workstation software-Quantitative Analysis.

### Cytochrome b_H_ Measurement

The method of cytochrome b_H_ measurement was adapted from that described by Quinlan et al. (61). Isolated mitochondria (1 mg/ml) were resuspended at 37 °C in buffer containing 120 mM KCl, 5 mM HEPES, 1 mM EGTA (pH 7.2 at 20 °C), and 0.3% (w/v) bovine serum albumin.

Absorbance change in Cytochrome b_H_ was measured after ODN or antimycin A addition using a NanoDrop One UV-Vis spectrophotometer (ThermoFisher Scientific). The Cytochrome b_H_ signal was recorded by spectrum scanning at the wavelength pair 566–575 nm at 37 °C with stirring and normalized based on the assumption that reductions of b_H_ are 0% with no added substrate and 100% with saturating substrates plus 2 µM DTT. In parallel with cytochrome b_H_ measurement, mtROS generation was detected in isolated mitochondria (0.5 mg/ml) treated by ODN or antimycin A using mitoSOX red as above described. To determine the effect of the mitochondria membrane potential on Cytochrome b_H_ redox state and mtROS generation in isolated mitochondria, b_H_ ^%red^ and mtROS were analyzed after addition of 500 nM FCCP in the assay buffer along with ODN or antimycin A treatment.

### Proteomics Analysis

The biotinylated ODN-bound proteins were precipitated with streptavidin beads from mitochondria lysates and resolved by PAGE (Bio-rad). The PAGE gels were stained using a silver staining kit (Pierce). Silver-stained gel pieces were excised, washed, destained and digested in-gel with 200 ng modified trypsin (sequencing grade, Promega) and Rapigest (TM, Waters Corp.) for 18 h at 37° C. In-solution samples were precipitated with 5:1 v/v of cold acetone at -20° C for 18 h, then centrifuged and the acetone was removed prior to treatment with Rapigest (100 °C for 10 min), followed by addition of trypsin. The resulting peptides were extracted and analyzed by high-sensitivity LC-MS/MS on an Orbitrap Fusion mass spectrometer (Thermo Scientific, Waltham MA). Proteins were identified by database searching of the fragment spectra against the SwissProt (EBI) protein database using Mascot (v 2.6, Matrix Science, London, UK) and Proteome Discoverer (v 2.2, Thermo Scientific). Typical search settings were: mass tolerances, 10 ppm precursor, 0.8d fragments; variable modifications, methionine sulfoxide, pyro-glutamate formation; enzyme, trypsin, up to 2 missed cleavages.

Peptides were subject to 1% FDR using reverse-database searching.

### RPPA Analysis

The reverse phase protein array (RPPA) analyses were performed to examine 161 protein targets in PBS or ODN treated HBEC3-KT cells. Cell lysates were serially diluted in 5 two-fold dilutions with RPPA lysis buffer. An Aushon Biosystems 2470 arrayer (Burlington, MA) was used to print lysates on nitrocellulose-coated FAST slides (Schleicher & Schuell BioScience) and make protein arrays. The array slides were probed with primary antibodies followed by detection with appropriate biotinylated secondary antibodies. The signal was amplified using the Vectastain ABC Elite kit (Vector Laboratories) and visualized by DAB colorimetric reaction. The slides were scanned, analyzed, and quantified using Microvigene software (VigeneTech Inc., Carlisle, MA) to generate spot signal intensities, which were processed by the R package SuperCurve (version 1.01). A fitted “supercurve” was plotted with the signal intensities on the Y-axis and the relative log2 concentration of each protein on the X-axis using the non-parametric, monotone increasing B-spline model (84). Protein concentrations were derived from the supercurve for each lysate by curve-fitting and normalized by median polish (85). Differential protein expression analysis was performed using R with LIMMA package and adjusted for multiple-testing using the Benjamini-Hoechberg method to reduce the false discovery rate.

### Quantification and Statistical Analysis

Statistical analyses were performed using SigmaPlot 14.0 (Systat Software, San Jose, CA) and GraphPad Prism 8 (GraphPad Software, San Diego, CA). One-way ANOVA was used to compare the means of multiple treatment conditions or multiple time points. The Holm-Sidak method was used, unless normality testing failed, in which case Kruskall-Wallis method was used. Means of two groups were compared using two-way Student’s t-test. Survival comparisons were performed using logrank testing by the Mantel-Cox approach.

## Extended Data and Supplementary Information

Tables S1 to S2

Fig. S1 to S15

## Acknowledgements

Supported by U.S. National Institutes of Health grants R01 HL117976, DP2 HL123229, and R35 HL144805 to S.E.E. and grants S10 OD012304-01 and P30 CA016672 to the Metabolomics Core Facility.

## Data and materials availability

The datasets generated for the Reverse-phase protein array data is available in the Evans Laboratory GitHub repository (www.github.com/evanslaboratory/Datasource). Further data information is available upon reasonable requests and should be directed to and will be fulfilled by the corresponding author, Scott E. Evans.

## References

1. Mizgerd JP. Lung infection--a public health priority. PLoS Med. 2006;3(2):e76.

2. File TM. Community-acquired pneumonia. Lancet. 2003;362(9400):1991–2001.

3. DALYs GBD, Collaborators H. Global, regional, and national disability-adjusted life-years (DALYs) for 333 diseases and injuries and healthy life expectancy (HALE) for 195 countries and territories, 1990-2016: a systematic analysis for the Global Burden of Disease Study 2016. Lancet. 2017;390(10100):1260–344.

4. Bjerre LM, Verheij TJ, Kochen MM. Antibiotics for community acquired pneumonia in adult outpatients. Cochrane Database Syst Rev. 2009(4):CD002109.

5. Niederman MS, Mandell LA, Anzueto A, Bass JB, Broughton WA, Campbell GD, et al. Guidelines for the management of adults with community-acquired pneumonia. Diagnosis, assessment of severity, antimicrobial therapy, and prevention. Am J Respir Crit Care Med. 2001;163(7):1730–54.

6. Leiva-Juarez MM, Kolls JK, Evans SE. Lung epithelial cells: therapeutically inducible effectors of antimicrobial defense. Mucosal Immunol. 2018;11(1):21–34.

7. Evans SE, Xu Y, Tuvim MJ, Dickey BF. Inducible innate resistance of lung epithelium to infection. Annu Rev Physiol. 2010;72:413–35.

8. Cleaver JO, You D, Michaud DR, Pruneda FA, Juarez MM, Zhang J, et al. Lung epithelial cells are essential effectors of inducible resistance to pneumonia. Mucosal Immunol. 2014;7(1):78–88.

9. Bals R, Hiemstra PS. Innate immunity in the lung: how epithelial cells fight against respiratory pathogens. Eur Respir J. 2004;23(2):327–33.

10. Bartlett JA, Fischer AJ, McCray PBJ. Innate immune functions of the airway epithelium. Contrib Microbiol. 2008;15:147–63.

11. Johnston SL, Goldblatt DL, Evans SE, Tuvim MJ, Dickey BF. Airway Epithelial Innate Immunity. Front Physiol. 2021;12:749077.

12. Clement CG, Evans SE, Evans CM, Hawke D, Kobayashi R, Reynolds PR, et al. Stimulation of lung innate immunity protects against lethal pneumococcal pneumonia in mice. Am J Respir Crit Care Med. 2008;177(12):1322–30.

13. Evans SE, Scott BL, Clement CG, Larson DT, Kontoyiannis D, Lewis RE, et al. Stimulated innate resistance of lung epithelium protects mice broadly against bacteria and fungi. Am J Respir Cell Mol Biol. 2010;42(1):40–50.

14. Tuvim MJ, Gilbert BE, Dickey BF, Evans SE. Synergistic TLR2/6 and TLR9 activation protects mice against lethal influenza pneumonia. PLoS One. 2012;7(1):e30596.

15. Duggan JM, You D, Cleaver JO, Larson DT, Garza RJ, Guzman Pruneda FA, et al. Synergistic interactions of TLR2/6 and TLR9 induce a high level of resistance to lung infection in mice. J Immunol. 2011;186(10):5916–26.

16. Evans SE, Tseng CK, Scott BL, Hook AM, Dickey BF. Inducible Epithelial Resistance against Coronavirus Pneumonia in Mice. Am J Respir Cell Mol Biol. 2020;63(4):540–1.

17. Kirkpatrick CT, Wang Y, Leiva Juarez MM, Shivshankar P, Pantaleon Garcia J, Plumer AK, et al. Inducible Lung Epithelial Resistance Requires Multisource Reactive Oxygen Species Generation To Protect against Viral Infections. mBio. 2018;9(3).

18. Ware HH, Kulkarni VV, Wang Y, Pantaleon Garcia J, Leiva Juarez M, Kirkpatrick CT, et al. Inducible lung epithelial resistance requires multisource reactive oxygen species generation to protect against bacterial infections. PLoS One. 2019;14(2):e0208216.

19. Galvan DL, Badal SS, Long J, Chang BH, Schumacker PT, Overbeek PA, et al. Real-time in vivo mitochondrial redox assessment confirms enhanced mitochondrial reactive oxygen species in diabetic nephropathy. Kidney Int. 2017;92(5):1282–7.

20. Guzman JN, Sanchez-Padilla J, Wokosin D, Kondapalli J, Ilijic E, Schumacker PT, et al. Oxidant stress evoked by pacemaking in dopaminergic neurons is attenuated by DJ-1. Nature. 2010;468(7324):696–700.

21. Wilson DF, Harrison DK, Vinogradov SA. Oxygen, pH, and mitochondrial oxidative phosphorylation. J Appl Physiol (1985). 2012;113(12):1838–45.

22. Zhao RZ, Jiang S, Zhang L, Yu ZB. Mitochondrial electron transport chain, ROS generation and uncoupling (Review). Int J Mol Med. 2019;44(1):3–15.

23. Murphy MP. How mitochondria produce reactive oxygen species. Biochem J. 2009;417(1):1–13.

24. Krieg AM. CpG motifs in bacterial DNA and their immune effects. Annu Rev Immunol. 2002;20:709–60.

25. Camara AKS, Zhou Y, Wen PC, Tajkhorshid E, Kwok WM. Mitochondrial VDAC1: A Key Gatekeeper as Potential Therapeutic Target. Front Physiol. 2017;8:460.

26. Vyssokikh MY, Brdiczka D. The function of complexes between the outer mitochondrial membrane pore (VDAC) and the adenine nucleotide translocase in regulation of energy metabolism and apoptosis. Acta Biochim Pol. 2003;50(2):389–404.

27. DeHart DN, Fang D, Heslop K, Li L, Lemasters JJ, Maldonado EN. Opening of voltage dependent anion channels promotes reactive oxygen species generation, mitochondrial dysfunction and cell death in cancer cells. Biochem Pharmacol. 2018;148:155–62.

28. Huo H, Zhou Z, Qin J, Liu W, Wang B, Gu Y. Erastin Disrupts Mitochondrial Permeability Transition Pore (mPTP) and Induces Apoptotic Death of Colorectal Cancer Cells. PLoS One. 2016;11(5):e0154605.

29. Cadenas S, Buckingham JA, St-Pierre J, Dickinson K, Jones RB, Brand MD. AMP decreases the efficiency of skeletal-muscle mitochondria. Biochem J. 2000;351 Pt 2:307–11.

30. Haworth RA, Hunter DR. Control of the mitochondrial permeability transition pore by high-affinity ADP binding at the ADP/ATP translocase in permeabilized mitochondria. J Bioenerg Biomembr. 2000;32(1):91–6.

31. Karch J, Bround MJ, Khalil H, Sargent MA, Latchman N, Terada N, et al. Inhibition of mitochondrial permeability transition by deletion of the ANT family and CypD. Sci Adv. 2019;5(8):eaaw4597.

32. Lemasters JJ. Evolution of Voltage-Dependent Anion Channel Function: From Molecular Sieve to Governator to Actuator of Ferroptosis. Front Oncol. 2017;7:303.

33. Carling D, Mayer FV, Sanders MJ, Gamblin SJ. AMP-activated protein kinase: nature’s energy sensor. Nat Chem Biol. 2011;7(8):512–8.

34. Hardie DG. AMP-activated/SNF1 protein kinases: conserved guardians of cellular energy. Nat Rev Mol Cell Biol. 2007;8(10):774–85.

35. Marcinko K, Steinberg GR. The role of AMPK in controlling metabolism and mitochondrial biogenesis during exercise. Exp Physiol. 2014;99(12):1581–5.

36. Steinberg GR. Role of the AMP-activated protein kinase in regulating fatty acid metabolism during exercise. Appl Physiol Nutr Metab. 2009;34(3):315–22.

37. Schlaepfer IR, Joshi M. CPT1A-mediated Fat Oxidation, Mechanisms, and Therapeutic Potential. Endocrinology. 2020;161(2).

38. Thupari JN, Pinn ML, Kuhajda FP. Fatty acid synthase inhibition in human breast cancer cells leads to malonyl-CoA-induced inhibition of fatty acid oxidation and cytotoxicity. Biochem Biophys Res Commun. 2001;285(2):217–23.

39. Wang Y, Palmfeldt J, Gregersen N, Makhov AM, Conway JF, Wang M, et al. Mitochondrial fatty acid oxidation and the electron transport chain comprise a multifunctional mitochondrial protein complex. J Biol Chem. 2019;294(33):12380–91.

40. Chowdhury SK, Gemin A, Singh G. High activity of mitochondrial glycerophosphate dehydrogenase and glycerophosphate-dependent ROS production in prostate cancer cell lines. Biochem Biophys Res Commun. 2005;333(4):1139–45.

41. Fendt SM, Bell EL, Keibler MA, Olenchock BA, Mayers JR, Wasylenko TM, et al. Reductive glutamine metabolism is a function of the alpha-ketoglutarate to citrate ratio in cells. Nat Commun. 2013;4:2236.

42. Huang LS, Shen JT, Wang AC, Berry EA. Crystallographic studies of the binding of ligands to the dicarboxylate site of Complex II, and the identity of the ligand in the “oxaloacetate-inhibited” state. Biochim Biophys Acta. 2006;1757(9-10):1073–83.

43. Kolaj-Robin O, O’Kane SR, Nitschke W, Leger C, Baymann F, Soulimane T. Biochemical and biophysical characterization of succinate: quinone reductase from Thermus thermophilus. Biochim Biophys Acta. 2011;1807(1):68–79.

44. Guaras A, Perales-Clemente E, Calvo E, Acin-Perez R, Loureiro-Lopez M, Pujol C, et al. The CoQH2/CoQ Ratio Serves as a Sensor of Respiratory Chain Efficiency. Cell Rep. 2016;15(1):197–209.

45. Brand MD. The sites and topology of mitochondrial superoxide production. Exp Gerontol. 2010;45(7-8):466–72.

46. Koopman WJ, Nijtmans LG, Dieteren CE, Roestenberg P, Valsecchi F, Smeitink JA, et al. Mammalian mitochondrial complex I: biogenesis, regulation, and reactive oxygen species generation. Antioxid Redox Signal. 2010;12(12):1431–70.

47. Li Y, Trush MA. Diphenyleneiodonium, an NAD(P)H oxidase inhibitor, also potently inhibits mitochondrial reactive oxygen species production. Biochem Biophys Res Commun. 1998;253(2):295–9.

48. Lambert AJ, Brand MD. Inhibitors of the quinone-binding site allow rapid superoxide production from mitochondrial NADH:ubiquinone oxidoreductase (complex I). J Biol Chem. 2004;279(38):39414–20.

49. Crofts AR, Hong S, Ugulava N, Barquera B, Gennis R, Guergova-Kuras M, et al. Pathways for proton release during ubihydroquinone oxidation by the bc(1) complex. Proc Natl Acad Sci U S A. 1999;96(18):10021–6.

50. Trumpower BL. The protonmotive Q cycle. Energy transduction by coupling of proton translocation to electron transfer by the cytochrome bc1 complex. xsJ Biol Chem. 1990;265(20):11409–12.

51. Lanciano P, Khalfaoui-Hassani B, Selamoglu N, Ghelli A, Rugolo M, Daldal F. Molecular mechanisms of superoxide production by complex III: a bacterial versus human mitochondrial comparative case study. Biochim Biophys Acta. 2013;1827(11-12):1332–9.

52. Lee I, Bender E, Arnold S, Kadenbach B. New control of mitochondrial membrane potential and ROS formation--a hypothesis. Biol Chem. 2001;382(12):1629–36.

53. Mazat JP, Devin A, Ransac S. Modelling mitochondrial ROS production by the respiratory chain. Cell Mol Life Sci. 2020;77(3):455–65.

54. Iwasaki A, Medzhitov R. Toll-like receptor control of the adaptive immune responses. Nat Immunol. 2004;5(10):987–95.

55. Klinman DM. Immunotherapeutic uses of CpG oligodeoxynucleotides. Nat Rev Immunol. 2004;4(4):249–58.

56. Kwon HJ, Lee KW, Yu SH, Han JH, Kim DS. NF-kappaB-dependent regulation of tumor necrosis factor-alpha gene expression by CpG-oligodeoxynucleotides. Biochem Biophys Res Commun. 2003;311(1):129–38.

57. Lee KW, Kim DS, Kwon HJ. CG sequence- and phosphorothioate backbone modification-dependent activation of the NF-kappaB-responsive gene expression by CpG-oligodeoxynucleotides in human RPMI 8226 B cells. Mol Immunol. 2004;41(10):955–64.

58. Suwarti S, Yamazaki T, Svetlana C, Hanagata N. Recognition of CpG oligodeoxynucleotides by human Toll-like receptor 9 and subsequent cytokine induction. Biochem Biophys Res Commun. 2013;430(4):1234–9.

59. Brandt U. Bifurcated ubihydroquinone oxidation in the cytochrome bc1 complex by proton-gated charge transfer. Febs Lett. 1996;387(1):1–6.

60. Hunte C, Palsdottir H, Trumpower BL. Protonmotive pathways and mechanisms in the cytochrome bc1 complex. Febs Lett. 2003;545(1):39–46.

61. Quinlan CL, Gerencser AA, Treberg JR, Brand MD. The mechanism of superoxide production by the antimycin-inhibited mitochondrial Q-cycle. J Biol Chem. 2011;286(36):31361–72.

62. Sun J, Trumpower BL. Superoxide anion generation by the cytochrome bc1 complex. Arch Biochem Biophys. 2003;419(2):198–206.

63. Rottenberg H, Covian R, Trumpower BL. Membrane potential greatly enhances superoxide generation by the cytochrome bc1 complex reconstituted into phospholipid vesicles. J Biol Chem. 2009;284(29):19203–10.

64. Federico A, Cardaioli E, Da Pozzo P, Formichi P, Gallus GN, Radi E. Mitochondria, oxidative stress and neurodegeneration. J Neurol Sci. 2012;322(1-2):254–62.

65. Minelli A, Bellezza I, Conte C, Culig Z. Oxidative stress-related aging: A role for prostate cancer? Biochim Biophys Acta. 2009;1795(2):83–91.

66. Asami DK, McDonald RB, Hagopian K, Horwitz BA, Warman D, Hsiao A, et al. Effect of aging, caloric restriction, and uncoupling protein 3 (UCP3) on mitochondrial proton leak in mice. Exp Gerontol. 2008;43(12):1069–76.

67. Lapointe J, Hekimi S. Early mitochondrial dysfunction in long-lived Mclk1+/-mice. J Biol Chem. 2008;283(38):26217–27.

68. Lee SJ, Hwang AB, Kenyon C. Inhibition of respiration extends C. elegans life span via reactive oxygen species that increase HIF-1 activity. Curr Biol. 2010;20(23):2131–6.

69. Scialo F, Sriram A, Fernandez-Ayala D, Gubina N, Lohmus M, Nelson G, et al. Mitochondrial ROS Produced via Reverse Electron Transport Extend Animal Lifespan. Cell Metab. 2016;23(4):725–34.

70. Yee C, Yang W, Hekimi S. The intrinsic apoptosis pathway mediates the pro-longevity response to mitochondrial ROS in C. elegans. Cell. 2014;157(4):897–909.

71. Bai Y, Onuma H, Bai X, Medvedev AV, Misukonis M, Weinberg JB, et al. Persistent nuclear factor-kappa B activation in Ucp2-/-mice leads to enhanced nitric oxide and inflammatory cytokine production. J Biol Chem. 2005;280(19):19062–9.

72. Wang D, Malo D, Hekimi S. Elevated mitochondrial reactive oxygen species generation affects the immune response via hypoxia-inducible factor-1alpha in long-lived Mclk1+/-mouse mutants. J Immunol. 2010;184(2):582–90.

73. Wang D, Wang Y, Argyriou C, Carriere A, Malo D, Hekimi S. An enhanced immune response of Mclk1(+)/(-) mutant mice is associated with partial protection from fibrosis, cancer and the development of biomarkers of aging. PLoS One. 2012;7(11):e49606.

74. Han D, Antunes F, Canali R, Rettori D, Cadenas E. Voltage-dependent anion channels control the release of the superoxide anion from mitochondria to cytosol. J Biol Chem. 2003;278(8):5557–63.

75. Muller FL, Liu Y, Van Remmen H. Complex III releases superoxide to both sides of the inner mitochondrial membrane. J Biol Chem. 2004;279(47):49064–73.

76. You Y, Richer EJ, Huang T, Brody SL. Growth and differentiation of mouse tracheal epithelial cells: selection of a proliferative population. Am J Physiol Lung Cell Mol Physiol. 2002;283(6):L1315–21.

77. Hemmi H, Takeuchi O, Kawai T, Kaisho T, Sato S, Sanjo H, et al. A Toll-like receptor recognizes bacterial DNA. Nature. 2000;408(6813):740–5.

78. Okubo T, Knoepfler PS, Eisenman RN, Hogan BL. Nmyc plays an essential role during lung development as a dosage-sensitive regulator of progenitor cell proliferation and differentiation. Development. 2005;132(6):1363–74.

79. Fasbender A, Lee JH, Walters RW, Moninger TO, Zabner J, Welsh MJ. Incorporation of adenovirus in calcium phosphate precipitates enhances gene transfer to airway epithelia in vitro and in vivo. J Clin Invest. 1998;102(1):184–93.

80. Cordero-Reyes AM, Gupte AA, Youker KA, Loebe M, Hsueh WA, Torre-Amione G, et al. Freshly isolated mitochondria from failing human hearts exhibit preserved respiratory function. J Mol Cell Cardiol. 2014;68:98–105.

81. Brautbar A, Wang J, Abdenur JE, Chang RC, Thomas JA, Grebe TA, et al. The mitochondrial 13513G>A mutation is associated with Leigh disease phenotypes independent of complex I deficiency in muscle. Mol Genet Metab. 2008;94(4):485–90.

82. Enns GM, Hoppel CL, DeArmond SJ, Schelley S, Bass N, Weisiger K, et al. Relationship of primary mitochondrial respiratory chain dysfunction to fiber type abnormalities in skeletal muscle. Clin Genet. 2005;68(4):337–48.

83. Vu TH, Sciacco M, Tanji K, Nichter C, Bonilla E, Chatkupt S, et al. Clinical manifestations of mitochondrial DNA depletion. Neurology. 1998;50(6):1783–90.

84. Tibes R, Qiu Y, Lu Y, Hennessy B, Andreeff M, Mills GB, et al. Reverse phase protein array: validation of a novel proteomic technology and utility for analysis of primary leukemia specimens and hematopoietic stem cells. Mol Cancer Ther. 2006;5(10):2512–21.

85. Hu J, He X, Baggerly KA, Coombes KR, Hennessy BT, Mills GB. Non-parametric quantification of protein lysate arrays. Bioinformatics. 2007;23(15):1986–94.

